# Single-molecule flow cytometry

**DOI:** 10.1101/2025.08.26.672174

**Authors:** Amir Rahmani, Matthew Christie, Amy Truesdale, James Thorne, Aleks Ponjavic

## Abstract

Flow cytometry (FC) is a powerful tool for high-throughput characterisation of cell populations, yet its ensemble fluorescence detection and the autofluorescence of cells impose a sensitivity limit of ∼100-1,000 fluorophores per cell. This precludes detection of low-abundance proteins and weak signalling events with biological importance and therapeutic potential. We introduce single-molecule Flow Cytometry (smFC), integrating high-numerical aperture oblique plane microscopy (OPM) with microfluidics to achieve optical sectioning and photon collection efficiencies suitable for single-molecule detection on flowing cells. Using super-bright, large Stokes shift fluorophores, smFC can detect labelled membrane proteins on cells with digital precision down to ∼2 molecules per cell, compared to an unlabelled control. This represents an improvement in the detection limit of 10- to 80-fold depending on the probe. We apply smFC to quantify the distribution of the membrane receptor c-kit in triple-negative breast cancer cells with unprecedented sensitivity. This revealed a previously unidentified distribution of 60% c-kit negative cells and 40% with low-abundance but heterogenous (1-200 mol/cell) surface distribution that is undetectable by conventional FC. These results establish smFC as a robust platform for high-throughput, quantitative single-molecule analysis in cell populations, opening new avenues for detecting rare biomarkers and quantifying the presence of low-abundance membrane proteins and targetable surface molecules with digital precision.

## Introduction

Flow cytometry (FC) has long been a cornerstone of cell analysis, enabling high-throughput quantification of membrane protein expression and isolation of phenotypes across large populations. ^1–5^ However, despite its ubiquity, FC fundamentally lacks the sensitivity to detect individual fluorescent molecules on the surface of cells, precluding the identification of proteins with low expression levels, weak signalling events, or heterogeneous expression patterns across cell populations. ^6^ Conventional FCs typically use single-pixel detectors that integrate fluorescence over the entire cell volume, making it impossible to distinguish dim signals from cellular autofluorescence. This sets a practical detection limit, typically around 100^7^ to 1,000^8^ fluorophores per cell, depending on the fluorophore and cell type. In most studies, a threshold is typically set to quantify the percentage of positive cells or the existence of a particular antigen. The outcome is that if a key molecule exists at low density, e.g. a receptor on an immune cell ^9^ or a targetable antigen on a tumour cell, ^10^ it will not be detected using FC. Therefore, bringing single-molecule sensitivity in cells to FC would allow identification of low-abundance and potentially unknown proteins and targetable antigens that are involved in vital signalling processes and disease.

Several studies have demonstrated the power of sensitive FC measurements on small particles such as extracellular vesicles, ^11,12^ virus-like particles, ^13^ or functionalised liposomes and exosomes. ^14^ While FC can achieve single-molecule sensitivity when studying these small objects, they lack the optical and biological complexity of cells. A recently developed digital flow cytometer (dFC) ^15^ achieved single-molecule detection of antibody-dye conjugates in vitro, but its reliance on ensemble signal integration and bead-based samples makes it unsuitable for resolving molecules on or within cells.

Imaging Flow Cytometry (IFC) ^16^ emerged to add morphological information and improved sensitivity by integrating imaging with flow-based measurements. However, existing IFC systems operate at relatively low NA, whereas high NA is typically required for single-molecule imaging to maximise photon collection prior to photobleaching. ^17^ Commercial IFC also uses widefield illumination, which does not provide optical sectioning that is essential for rejecting out-of-focus fluorescence background to maximise contrast. ^18^ Light-sheet illumination has recently been applied to IFC to overcome this limitation; ^19–21^ however, these implementations either suffer from limited spatial resolution and low photon collection efficiency due to low NA or are incompatible with standard microfluidic mounting geometries. As a result, they are not readily adaptable for applications that require high detection sensitivity such as single-molecule counting.

Given the promise of improved detection in FC, a number of avenues have been explored to overcome the sensitivity limit. ^8^ Rolling circle amplification using biotin-labelled antibodies provides a fluorescence signal boost that makes single molecules visible using conventional epifluorescence imaging. ^22^ However this is a slow and arduous process that is also only compatible with fixed cells, preventing sorting and downstream analysis. Ultrabright multimeric streptavidin-phycoerythrin (SA-PE) conjugates enable epifluorescence detection of individual antibodies, ^23^ however this has not been used with FC and the influence and biocompatibility of these multimeric particles is unknown. Single-molecule lo-calisation microscopy has also been used to demonstrate superior sensitivity to FC. ^24^ This approach solved a problem in anti-CD19 immunotherapy for leukaemia, where it was shown that anti-CD19 treatment was effective even for patients that were determined CD19^−^ according to FC. Super-resolution imaging revealed that these CD19^−^ cells actually had a small number CD19 that was sufficient for treatment but could not be detected by FC. Of course, such methodology is incompatible with FC due to special buffers and slow imaging, but this demonstrates the potential of enabling single-molecule sensitivity in FC.

Here, we introduce single-molecule Flow Cytometry (smFC), based on high-NA oblique plane microscopy (OPM), ^25–27^ integrated with a microfluidic flow system. By combining the optical sectioning capabilities of OPM with recently available superbright fluorophores, ^28^ smFC achieves the sensitivity and reproducibility required to detect and quantify individual molecules in flowing cells with digital accuracy. We show that for the same fluorophore, smFC affords a 50-fold improvement compared to FC in the limit of detection, down to single-molecule (1-2 molecules/cell) levels. Finally, we apply smFC to clarify the controversy surrounding the expression of the c-kit stem cell marker on the surface of triple-negative breast cancer (TNBC) cells. Several studies utilising conventional FC have reported MDA-MB-468 cells as c-kit negative, ^29,30^ while others report this TNBC cell line as c-kit positive. ^31^

## Results and Discussion

### Oblique plane microscopy for 3D counting of individual proteins on cells

A key requirement for achieving single-molecule sensitivity in FC is the ability to perform three-dimensional imaging as cells flow across the imaging plane. To address this, we implemented a high-NA (1.1 in shear-direction and 1.2 in perpendicular axis) OPM system based on an existing design. ^27^ OPM can image a tilted plane within the sample, which enables volumetric sampling as cells move horizontally through the imaging plane (Figure 1a). We introduced a 30° inclined light sheet, with a thickness of 2 *µ*m FWHM and a propagation length of 30 *µ*m (Figure S1), to cover a 25 *µ*m height, suitable for 3D cellular imaging. Although the tilted detection in OPM reduces fluorescence transmission (58.5% compared to epi-fluorescence, Figure S2), this is offset by the improved signal-to-background ratio (SNR) due to optical sectioning, which has been demonstrated in OPM super-resolution microscopy. ^32^

**Fig. 1.**
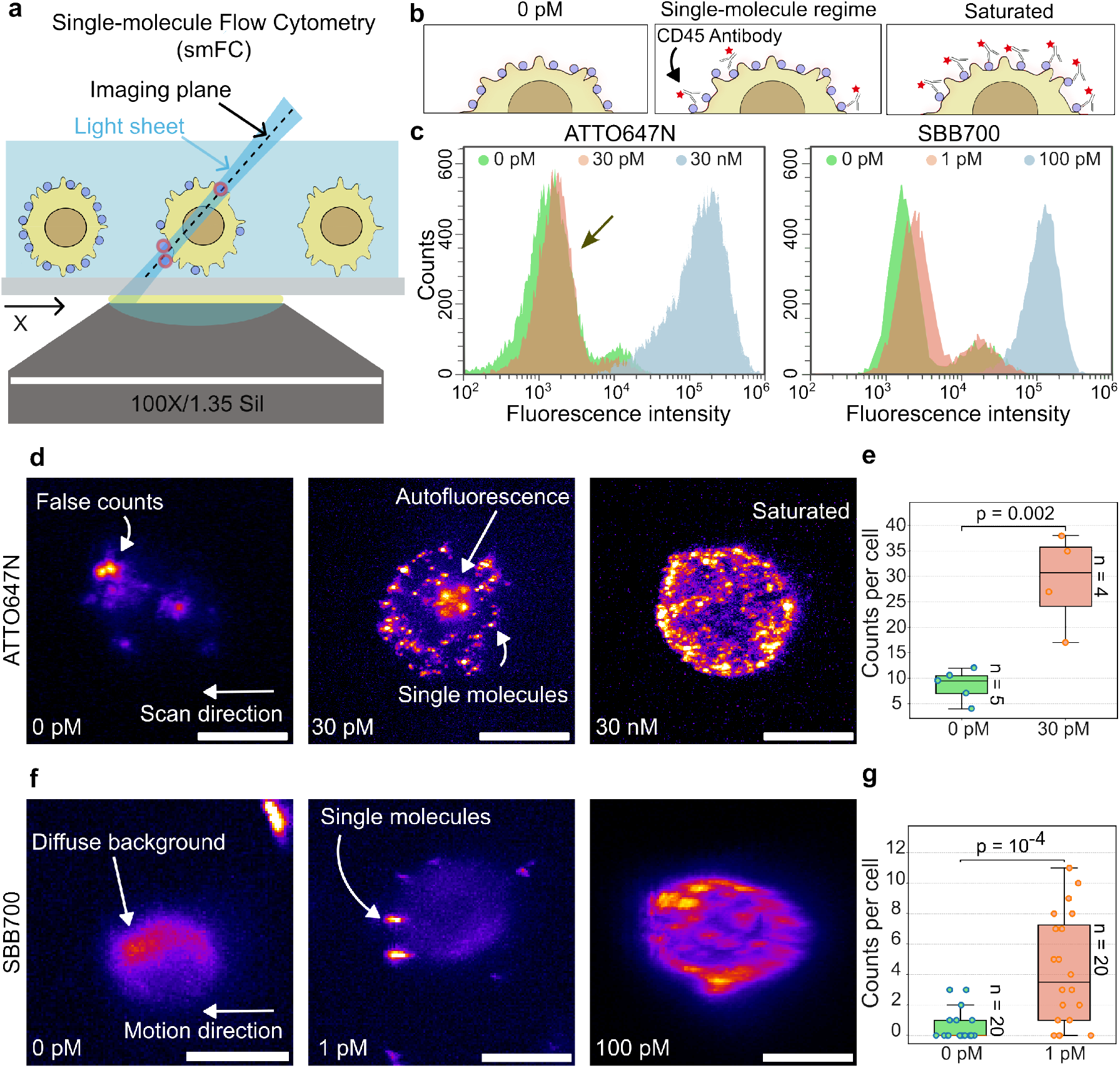
Single-molecule counting on cells using high-NA OPM microscopy. (a) Schematic of the smFC system integrating oblique light-sheet illumination/detection enabling volumetric imaging. (b) Conceptual illustration of membrane protein labelling on cells for three conditions: (1) 0 pM unlabelled control, (2) low-concentration labelling (single-molecule regime), and standard labelling (saturating antibody concentration). (c) Left: Flow cytometry histograms of ATTO647N/CD45-labelled fixed Jurkat T cells at three labelling levels: unlabelled (control), low concentration (3 pM), and standard concentration (30 nM). Only the standard-labelled sample shows a distinct fluorescence peak, while the low-labelled population overlaps with the control (indicated by arrow), highlighting the detection limit of conventional FC. Right: Similar for SBB700/CD45. (d) Maximum intensity projection images from a deskewed z-stack of ATTO647N/CD45-labelled Jurkat T cells. Cells were imaged in 3D using OPM by scanning with x-translation, revealing individual fluorophores localised on the membrane. Scale bar = 10 *µ*m. (e) ATTO647N counts using OPM for the control (0 pM) and single-molecule regime (30 pM). Means reported and error bars represent standard deviation. (f) as for d but with secondary SBB700/CD45 labelling and with the cells in continuous motion to emulate the effects of flow. (g) As for (e) but for SBB700/CD45.

High-speed single-molecule imaging relies on photostable fluorophores with high photon flux to ensure reliable detection within short exposure times (<5 ms). To evaluate our system for this purpose, we initially labelled Jurkat T cells using ATTO647N-conjugated primary antibodies. ATTO647N is commonly used in smFRET ^33^ and DNA-PAINT ^34^ applications due to its high photon rates and photostability, even at ms intervals. For standard labelling, we applied the recommended antibody concentration of 30 nM, while for achieving single-molecule labelling regime, we reduced the concentration to 30 pM to ensure sparse labelling. An overview of labelling is shown in Figure 1b. We targeted the phosphatase CD45 as it is a well-characterised membrane protein that is highly expressed on the stable Jurkat T cell line. ^35^

### Flow Cytometry requires >100-1,000 molecules/cells

We first carried out FC measurements using the recommended labelling concentration (30 nM) on a commercial Cytoflex S cytometer, which revealed abundant expression of CD45 on fixed Jurkat T cells (Figure 1c). Using a PE-labelled quantitation kit (Quantibrite beads) and PE anti-CD45 antibodies, we were able to determine the number of CD45 molecules on Jurkat T cells to be ~100,000 at 30 nM of primary antibody, similar to other reports. ^35^ However, when the concentration was lowered *∼*1,000-fold to 30 pM (expected ~100 mol/cell based on PE quantitation), the distribution was virtually identical to the negative control (Figure 1c), demonstrating the substantial presence of autofluorescence that was masking the signal from the low abundance of labelled antibodies. Based on the FC measurements we could determine a limit of detection ^36^ of *∼*2,000 mol/cell. While ATTO647N is ideal for single-molecule measurements, it is not commonly used in FC, perhaps because of this relatively poor detection limit, although the limit of FC has been quoted in ranges between *∼*100-1,000 mol/cell ^7,8^ depending on the fluorophore used. We also observed relatively substantial autofluorescence in the far-red region, inherent to Jurkat T cells. Note that because we use the geometric mean for FC analysis this should not affect the limit of detection as the geometric mean is robust to outliers.

### OPM improves the detection limit of FC 80-fold for ATTO647N

Next, we placed cells from the same aliquots measured using FC on our OPM-based smFC instrument and acquired 3D intensity distributions using a relatively high exposure time of 30 ms for quantification purposes. We initially imaged cells labelled with a relatively low concentration of ATTO647N/CD45 primary antibody (300 pM, 100× less than recommended). Despite the low concentration we observed bright membrane staining (Video S1), visualised as a deskewed z-stack max projection in Figure 1d. At this level, the distribution of CD45 is too dense to count individual molecules. However, when we imaged cells at low antibody concentration (30 pM, expected 30-60 mol/cell), for which FC could not detect anything, we observed clear point-like distributions, representative of individual molecules (29.3±9.4 mol/cell, Figure 1e). While this approach allowed us to quantify the presence of low copy numbers of CD45 on Jurkat T cells (p = 0.002), we also observed point-like patterns on unlabelled cells (8.8±3.3 mol/cell, Figure 1e, Video S1), due to the presence of autofluorescence. Based on the assumption that this originated from lipofuscin accumulation, we attempted to quench the autofluorescence using TrueBlack and TrueBlack Plus, but this was unsuccessful. Using ATTO647N we achieved a limit of detection of 29 mol/cell, representing an *∼*80-fold improvement compared to conventional FC. While impressive and useful, this approach would not achieve smFC with digital accuracy.

### Large Stokes shift probes enable digital counting using OPM

We considered the typical spectra of autofluorescence in cells and hypothesised that use of a bright probe with a large Stokes shift might limit the influence of autofluorescence in a red-shifted channel. We trialled several tandem probes (Brilliant Violet and RPE-Astral616) as well as conventional ultrabright probes (PE) for this purpose, and while these were all sufficiently bright for single-molecule detection with OPM, they photobleached far too quickly (<1 frame) to be useful. Ultimately, we found that a polymer dot-based probe, ^37^ Starbright Blue 700 (SBB700), offered both the required brightness and large Stokes shift, as well as superior photostability, to achieve our goals. We chose to use secondary labelling for compatibility with multiple protein targets. It should be noted that this maintains live-cell compatibility allowing FACS, and all our subsequent experiments were carried out with living cells. Furthermore, a primary antibody conjugation kit is in development for Starbright dyes and the probe is available as a primary label.

To assess the performance of the large Stokes shift probe, we imaged Jurkat T cells labelled with mouse anti-human CD45 primary antibodies, followed by secondary labelling with SBB700 goat anti-mouse IgG (Figure 1f). To prepare for future FC experiments, we now performed imaging with low exposure time (5 ms, 198 Hz frame rate), reduced resolution for speed (2×2 binning, 228 nm pixel size) and with cells in motion (4 pixels/frame), achieved by moving the microscope stage at a fixed velocity. At moderate primary antibody concentration (100 pM), we again observed substantial membrane staining (Figure 1f), while for an extremely low single-molecule concentration (1 pM, Video S2) we were able to detect 4.3±3.6 mol/cell (Figure 1g). The notable difference now was for the negative control (0.55±0.99 mol/cell). While background fluorescence from cellular autofluorescence was still present, the broader spectral separation allowed a greater proportion of the puncta to disappear with only diffuse background remaining (Figure 1f, Video S2). Furthermore, the intrinsic brightness of SBB700 ensured that detected emission events consistently produced high SNR values, without blinking, reducing the likelihood of false counts and enabling digital identification of as few as 2–3 molecules per cell with high confidence (p = 10^−4^). This combination of OPM-based optical sectioning and tailored fluorophore selection played a critical role in achieving the sensitivity and specificity necessary for robust single-molecule counting.

Naturally, the superior probe also improved the performance of FC for the cells from the same aliquot studied using OPM (Figure 1c). This now resulted in an impressive detection limit of 60 mol/cell for FC. Nevertheless, at the lowest concentration studied (1 pM), FC was unable to detect any substantial signal, while OPM could observe on average 4.3 mol/cell. Importantly, while the intrinsic background signal, which is primarily from cellular autofluorescence, is comparable between conventional FC and smFC, how this background interacts with the detection architecture differs significantly. In conventional FC, single-pixel detectors integrate photons across the entire illuminated volume, and the resulting signal is effectively averaged over the whole cell body. This reduces the signal-to-background ratio (SBR), which prevents detection of low-abundance fluorophores. In contrast, smFC uses a camera-based detection system with optical sectioning via oblique light-sheet illumination, which confines excitation to a thin plane and spatially localises emitted photons at the pixel level. This approach dramatically reduces the background per pixel by >100-fold, allowing bright fluorophores such as SBB700 to produce localised high-SNB spots that stand out clearly from autofluorescence. As a result, OPM can detect and count as few as 2–3 molecules per cell, a regime that is far below the practical detection limit of conventional FC.

### Flow cytometry with single-molecule sensitivity (smFC)

To achieve single-molecule sensitivity in a FC-compatible format, we integrated our OPM with a microfluidic platform designed for controlled lateral translation of cells through a thin light sheet (Figure 2a). The flow rate was set to achieve a mean cell velocity of 4 pixels/frame (*∼*200 µm/s), balancing the need for high-throughput operation with limited motion blur and sufficient exposure time (5 ms) to detect single emitters with adequate photon counts. The microfluidic channel height (25 *µ*m) was matched to the light sheet thickness (2 *µ*m) and Rayleigh length (30 *µ*m), ensuring consistent excitation of the cell membrane as cells passed through the field of view. We used a wide channel (300 *µ*m) to maximise cell throughput as the exposure time was fixed.

**Fig. 2.**
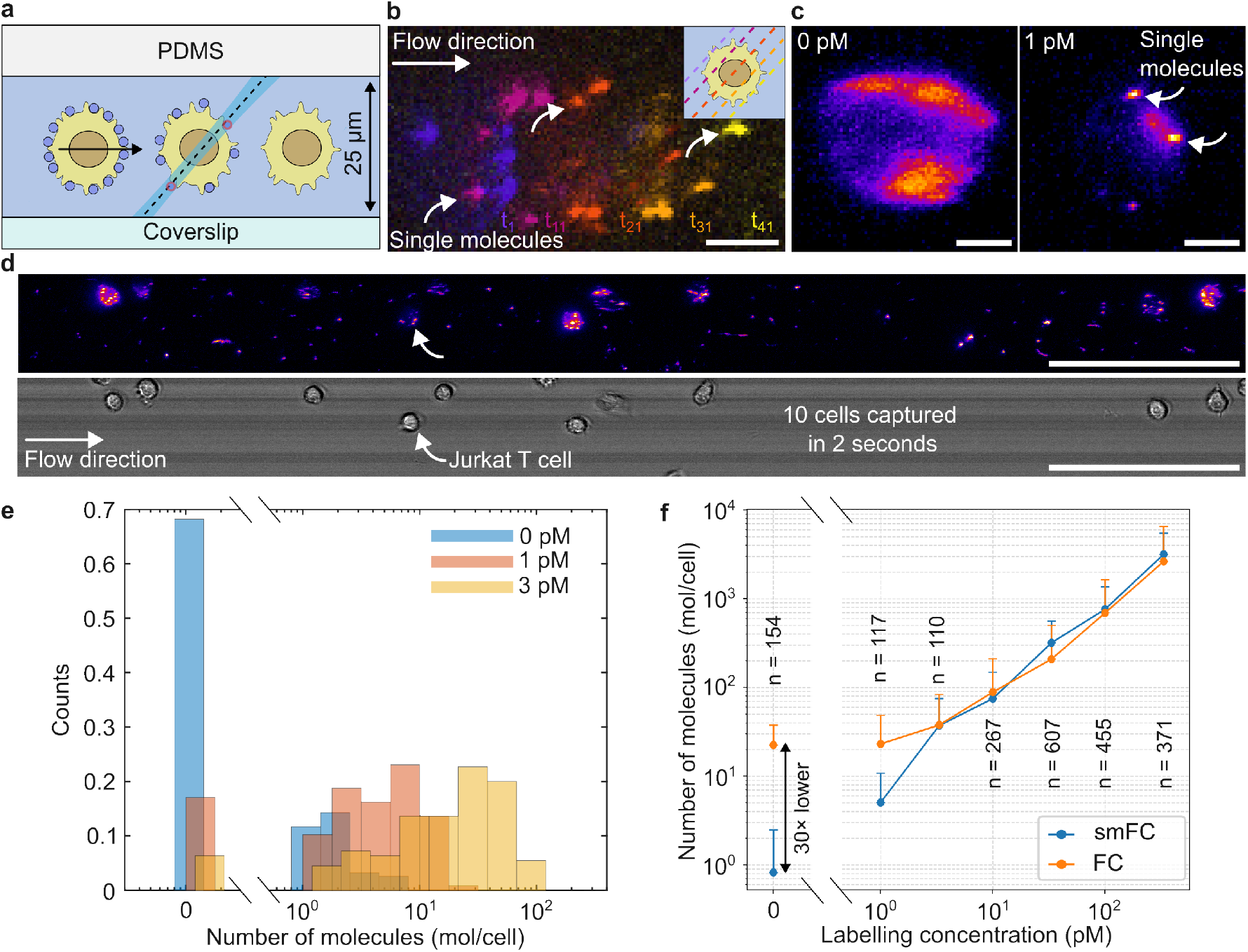
Quantitative single-molecule analysis of membrane proteins in flow using smFC. (a) Schematic of cells labelled with SBB700/CD45 in flow imaged with smFC. (b) Representative montage (t numbers correspond to frame) of a live Jurkat T cell labelled at low SBB700/CD45 primary antibody concentration (3 pM) imaged in flow using smFC. Individual membrane-bound fluorophores are visible. Scale bar = 5 *µ*m. (c) Maximum intensity projections of cells labelled at different concentrations imaged using smFC. Control (0 pM) shows diffuse background but no single molecules, while 1 pM shows three molecules. Scale bar = 5 *µ*m. (d) 2 seconds of smFC imaging in the fluorescence (top) and brightfield (bottom) channels, rapidly capturing data for 10 cells. Scale bar = 100 *µ*m. (e) CD45 distributions at different primary antibody concentrations, demonstrating sensitivity within the 1-10 mol/cell range. Bars are shifted for visual interpretation. (f) Comparison of quantification of cells from the same sample using smFC and conventional FC. Hybrid single-molecule/intensity analysis allows smFC to span the entire detection range (1-10,000 mol/cell), while affording a *∼*30 reduction in background signal for the control, allowing digital detection capability between 1-50 mol/cell. Data is presented as mean±standard deviation for smFC and geometric mean±robust standard deviation for FC to minimise effects of outliers.

We applied OPM imaging to quantify SBB700/CD45– labelled live Jurkat T cells (>10^6^ cells/ml) in flow, labelled across a range of primary antibody concentrations, from the single-molecule regime (1 pM) to approaching the saturated regime (300 pM). Continuous image acquisition was started once the cells were loaded into the channel. A timelapse of a cell labelled at 3 pM is shown in Figure 2b, demonstrating the ability of OPM/smFC to achieve 3D single-molecule coeunting using flow. Representative deskewed maximum z-projections of individual Jurkat T cells labelled at different SBB700/CD45 concentrations are shown in Figure 2c. At low primary labelling concentration (1 pM), we could resolve and count individual copies of CD45 on the cell membrane, while the control (0 pM) only displayed a diffuse background, similarly to the result for OPM imaging. A separate brightfield imaging channel was used to detect cells with no fluorescence signal and to ensure that fluorescence emission originated from labelled cells rather than debris or unbound antibodies (Figure 2d, Video S3). This channel was also used to confirm throughput, which ranged between 5-10 cells/s, compared to typical FC throughputs of 100-1,000 cells/s.

We first considered histograms of molecule counts per cell (Figure 2e) for the low primary antibody concentration regime (control, 1 pM, and 3 pM). These histograms, which represent the percentage of the cell population as a function of molecule counts per cell, revealed that for unlabelled cells (0 pM control), 70% of cells had a count of zero molecules and, importantly, 95% of cells had a count of 3 molecules or less. As the primary antibody concentration was increased, we measured on average 4.7±4.2 (17% zero) mol/cell and 21±18.4 (6% zero) for 1 and 3 pM respectively, demonstrating that the ability of OPM to count individual molecules on cells with digital precision was retained in a FC modality.

To quantify smFC sensitivity across a wide range of abundances (Video S4), we plotted the average number of molecules per cell as a function of primary antibody concentration (Figure 2f). Note that Figure 2f represents a hybrid analysis approach. At higher concentrations (>30 pM), the mol/cell value plateaued as individual receptors could no longer be identified due to overlapping PSFs, whereas the relationship remained proportional from 1 to 10 pM (Figure S4), consistent with the digital counting regime (*∼*3-50 mol/cell). A FC with such a limited dynamic range would offer limited use, which prompted us to consider for each cell the mean intensity as well as the localisation count. If cells had an intensity equivalent to 35 molecules or greater, the molecular count was estimated using intensity instead as done for conventional FC, allowing us to bridge the single-molecule and high-density regimes. A unique benefit of smFC is that as long as there are some cells in the single-molecule regime, this acts as an internal calibration for the intensity/receptor. Such a calibration could always be created by using low labelling concentrations as we have done here. These results demonstrate that our system can reliably count individual membrane-bound proteins on flowing cells under dynamic conditions, while maintaining a high dynamic range, enabling high-throughput single-molecule detection in heterogeneous cell populations.

### Quantifying the distribution of c-kit on triple-negative breast cancer cells

Having established the capabilities of smFC, we applied our highly sensitive cytometry platform to characterise a surface marker with biological and clinical relevance that elude conventional cytometry. We decided to focus on c-kit, a membrane protein that plays an important role in regulation of tumour formation and progression (Figure 3a). ^38^ This receptor recognises stem-cell factor (SCF) to stimulate cell growth and survival. Efforts have been made to target c-kit for anti-cancer immunetherapy in general, ^39^ but also specifically for triple-negative breast cancer (TNBC). ^31^ Despite its promise, the presence of c-kit on cell line models of TNBC is contentious. Here, we selected the TNBC cell line MDA-MB-468 that is commonly used in PD-1/PD-L1 blockade ^40^ and CAR T-cell ^41^ immunotherapy studies. Notably, some reports show near-detection limit expression of c-kit using flow cytometry on MDA-MB-468, ^31^ while others have found c-kit expression using proteomics ^42^ or immunoblots. ^43,44^ Others however state that MDA-MB-468 is negative for c-kit expression. ^29,30^ Application of smFC would therefore facilitate confident quantification of important markers on cell lines with clinical relevance.

**Fig. 3.**
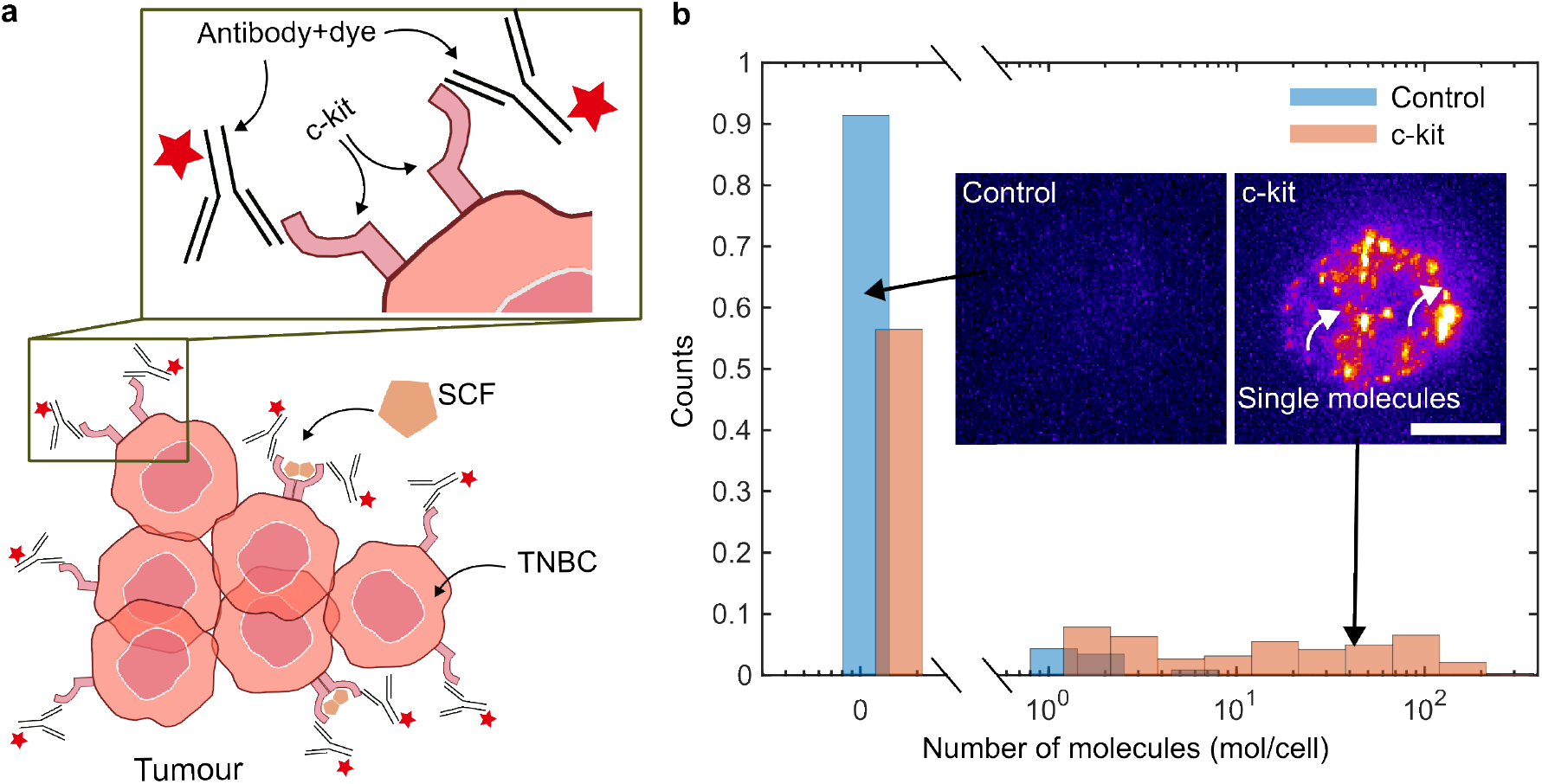
smFC enables quantification of low-abundance c-kit expression on TNBC cells. (a) Schematic illustration of a TNBC cell with c-kit expression on the surface. (b) smFC histogram showing c-kit detection on MDA-MB-468 cells, revealing a subpopulation with detectable surface expression. Bars are shifted for visual interpretation. The insets show representative images of a control (no primary) cell and SBB700/c-kit-labelled cells. Scale bar = 10 *µ*m.

Using smFC we found that 30% of MDA-MB-468 cells have more than 3 c-kit molecules on the cell surface (Figure 3b). For the control (no primary antibody), 90% had zero and 0.9% of cells had more than 3 false detection events. From the positive c-kit cells (above 3 molecules), the median number of molecules on the cell surface was 25, which is below the detection limit for conventional flow cytometers, even using SBB700 labelling. These results suggest that there is a wide distribution of TNBC cells with likely targetable but undetectable levels of c-kit, unless studied using smFC.

While previous work identified surface expression of c-kit on 85% of MDA-MB-468 cells, ^31^ this is based on overlap methods beyond the limit of detection, that while useful are prone to overinterpretation and detection of false events. ^45^ Furthermore, the result is in opposition to other work. ^29,30^ The detection limit of FC is likely a major cause for lack of consensus regarding the expression of c-kit on TNBC cell lines. Proteomics and western blots can identify the presence of proteins with high sensitivity, but these include intra-cellular proteins, and functional outcomes and therapeutics mostly on the presence of membrane proteins on the membrane. A highly sensitive method such as smFC is therefore required to improve characterisation of low-abundance membrane proteins. It should be noted that there are numerous low-expression cancer-related markers similar to c-kit, ^46^ such as CAR-T targets (CD19, BCMA, CD20 among others). ^47^ smFC would definitely confirm the presence or absence of these low-abundant, but therapeutically relevant targets on various cell line models of cancer and other diseases and offers the potential of identifying novel theranostic biomarkers.

Approximately 40-50% of TNBC/basal-like breast cancers are reported c-kit positive by IHC, ^48^ with the remainder considered negative. However, immunohistochemistry (IHC) and conventional FC are only capable of detection at the resolution of >100 mol/cell. smFC improves detection by 1-2 orders of magnitude and our analysis of a TNBC cell line identified three distinct subpopulations based on c-kit abundance (<1, 1–100, >100 molecules/cell). This enables stratification into true negative, low-expressing, and high abundance populations, with minimal false positives. Applying smFC to primary TNBC samples may therefore reveal previously hidden c-kit positive (albeit low abundance) tumours. Currently, c-kit positivity is not thought to associate with clinical outcome, ^49^ but improved quantification that can stratify into negative, low-, and high-expressing populations, coupled with improved mechanistic understanding, could prompt re-evaluation of these associations. Applying smFC to in vitro studies could transform understanding of the functional consequence of c-kit abundance and its mechanism of action. c-kit is targetable by FDA/NICE-approved TKIs such as Nilotinib and Imatinib for myeloid leukemia yet is not a therapeutic target in TNBC nor several other c-kit expressing cancers. The ability to distinguish truly negative c-kit tumours from low-expressing positive tumours opens new therapeutic options and avenues for understanding receptor-driven signalling, akin to EGFR pathways or T-cell activation, where even a single ligand-receptor interaction can trigger downstream effects. ^50,51^These studies are now in reach for many more pathways as smFC allows separation of truly negative cells from those with below IHC/convention FC detection limits.

### Advantages and limitations of smFC

smFC enables exploration of biological phenomena, currently inaccessible by conventional FC, such as studying heterogeneity in low-expression membrane receptors, rare signalling events, and phenotypically distinct subpopulations defined by low-abundance markers. This makes smFC suitable for quantifying targetable molecules on tumours or detecting sparse receptor engagement in drug assays. Furthermore, its utility can be extended to dynamic studies such as protein turnover or endocytosis, providing single-molecule sensitive data across statistically meaningful populations.

Future developments could integrate GPU-accelerated real-time single-molecule detection, ^52^ to enable on-the-fly gating and sorting decisions, advancing smFC towards a true digital cytometry and sorting platform. Throughput gains will require balancing of excitation time, motion blur, detection speed and importantly incorporation of flow focusing using sheath flow. Although there are faster detectors (e.g., Kinetix sCMOS or SPAD arrays) ^53^ the limiting factor remains the minimum excitation time (*∼*5 ms) to collect sufficient photons per molecule to be confident in an event. The signal could potentially be boosted through multi-label amplification (e.g., through clustered secondaries), though this compromises compatibility with live-cell assays. In terms of motion blur, it is possible to use scanning mirrors to counteract motion. ^54^ Because axial drift is a major challenge in OPM, ^55^ long-term performance of smFC can be improved by integrating an active focus stabilisation unit ^56^ to the microscope.

To match the multi-parameter capabilities of conventional FC, smFC can adapt hybrid operation: single-molecule counting in one or two primary channels, with conventional fluorescence readouts in additional channels for phenotyping. This approach would preserve digital accuracy for rare targets while enabling high-dimensional population classification, including co-localisation of extremely rare signalling events.

Ultimately, OPM is unlikely to be the most efficient modality for smFC due to the cost, complexity and limited fluorescence transmission of OPM. The throughput and accessibility could be improved by adopting single-molecule light-field microscopy that can capture the entire cell in one frame. ^57,58^ It should be noted that the upper level of countable molecules, as well as the sensitivity, would however likely be reduced due to the increased dimensions of the PSF and lack of optical sectioning.

While smFC can detect as few as 2-3 mol/cell compared to an unlabelled control, the utility of this depends on the performance of the primary antibody used and particularly its propensity for unspecific binding. We have shown that unspecific binding from the secondary SBB700 antibody is minimal under BSA-blocking conditions (Fig. 1g). To establish that the 1-10 mol/cell range is useful, we quantified a series of negative control antibodies (0.005 mg/ml, 30 nM saturating labelling concentration) on MDA-MB-231 (CD24^−^/c-kit^−^) TNBCs and on Jurkat T cells (CD8^−^), as well as a positive control antibody on MDA-MB-468 (CD24^+^) TNBCs using OPM. We observed essentially no events for negative antibodies on MDA-MB-231 (CD24: 0.56±1.3 mol/cell, c-kit: 0.89±0.95 mol/cell, no primary: 0.95±0.56 mol/cell, N.S. one-way ANOVA p = 0.42, Fig. S3) and on Jurkat T cells (CD8: 1.18±1.91 mol/cell, no primary: 0.60±1.13 mol/cell, N.S. p = 0.1, Fig. S3). On the other hand MDA-MB-468 was found to be positive for CD24 (CD24: 2967±1207 mol/cell, no primary: 0.31±0.62 mol/cell, p < 10-20, Fig. S3). This suggests that, given effective antibodies, smFC can truly detect the presence of a few molecules on the surface on cells.

## Conclusion

We have developed and characterised a high-NA OPM system optimised for single-molecule detection in both live and fixed cells in flow. Our system allows for 3D mapping and, notably, digital counting of individual molecules, even at an abundance of just 2–3 molecules per cell. Based on our fluorophore calibration and antibody-labelling efficiency, this corresponds to an improvement in the detection limit by 10- to 80-fold (depending on fluorophore) compared to commercial cytometers, which are typically limited to ≥100 molecules per cell.

smFC allows for exploring biological phenomena that are currently inaccessible using conventional flow cytometry. This can be applied to study heterogeneity in low-expression membrane receptors, to quantify rare signalling events, or to identify phenotypically distinct subpopulations defined by sparse expression patterns. The characterisation of c-kit in MDA-MB-468 highlights the ability of smFC to reveal the presence of membrane proteins or antigens with digital precision, which has the potential to: (1) Identify previously unknown or disputed low-abundance molecules on cells; (2) Improve the characterisation of cancer patient cells. In conclusion, smFC via OPM overcomes key sensitivity barriers of traditional platforms and opens new avenues in quantitative cell biology. Its integration into high-throughput workflows promises real-time, digital resolution of detecting rare molecular events, which can bring molecular-scale sensitivity into the domain of scalable cell analysis and sorting.

## Author Contributions

AR, and AP wrote the main manuscript. JT wrote sections on c-kit and TNBCs. The OPM system was implemented by AR under AP’s supervision. MC and AT designed and fabricated microfluidic devices and related protocols. AR and AP carried out all experiments. AR and AP performed data analysis. AR, JT and AP edited the manuscript. Visualization was carried out by AR and AP. All authors reviewed the manuscript.

## Acknowledgements

This work was funded by a University of Leeds University Academic Fellowship, a Royal Society Research Grant (RGS\ R2\ 202446), as well as an AMS Springboard Award (SBF006\1138), awarded to A.P., and by an EPSRC Bragg Centre Doctoral Training Studentship (EP/T517860/1) awarded to A.R. We would also like to acknowledge a Funding for Places award from The Wolfson Foundation, which has funded the Wolfson Imaging Facility at the University of Leeds, where the OPM/smFC is housed.

We would like to thank Ruth Hughes for advice regarding flow cytometry, Rosa Catania for assistance with dye-conjugation and purification, Graham Cook for correspondence on immunotherapy and flow cytometry, Michelle Peckham for donating the TrueBlack reagent, Benjamin Johnson for suggestions on microfluidics, Alistair Curd for managing/improving the OPM instrument, and, finally, Alfred Millet-Sikking, Reto Fiolka, James Manton, Brandon Scott and Andreas Bodén for assistance with trouble-shooting OPM alignment. Additionally, we would like to thank Bio-Rad representatives David Goldeck and Amelia Louver for helpful discussion and for providing product details as well as SBB700-conjugated primary antibody samples.

## Notes

The authors declare no competing financial interest.

## Methods

### Cell culture

Jurkat T cells (ATCC) were grown at 37°C with 5% CO_2_ in phenol red-free RPMI 1640 Medium (11835030, Sigma-Aldrich) supplemented with, 10 mM HEPES, 1 mM sodium pyruvate and 10% heat-inactivated FBS (F7524, Sigma-Aldrich).

TNBC (MDA-MB-468 and MDA-MB-231) cell lines (ATCC) were grown at 37°C with 5% CO_2_ in DMEM (31966047, Thermo Fisher) supplemented with 10% FBS (11560636, Thermo Fisher). TNBC cell lines were confirmed mycoplasma free at 6-monthly intervals, and cell line identities were authenticated at the start of the project.

### Antibodies

Cells were stained with purified mouse anti-human CD45 (Clone HI30, 304001, BioLegend), purified mouse anti-human CD117 (c-kit, Clone 104D2, 313201, BioLegend), purified mouse anti-human CD24 (Clone ML5, 311101, BioLegend), purified mouse anti-human CD8 (Clone SK1, 344702, BioLegend), PE anti-human CD45 (Clone 2D1, 368509, BioLegend), and secondary StarBright Blue 700 (SBB700) Goat Anti-Mouse IgG (12004159, Bio-Rad). Alternative probes were also evaluated including Brilliant Violet anti-human CD45 (Clone HI30, 304043, BioLegend) and RPE-Astral616 anti-human CD45 (Clone 2D1, P012-R616-125, Biotium).

### Primary antibody labelling

The NHS-ester of ATTO647N (AD 647N-31, ATTO-TEC) was reacted with anti-CD45 primary antibody. Sodium bicarbonate was added to 30 *µ*l of 1.5 mg/ml (concentrated using Vivaspin centrifugal concentrator) primary antibody in PBS to make up a concentration of 0.1 M sodium bicarbonate. NHS-ester of ATTO647N was then added at a final concentration of 100 *µ*M. The relatively high excess was required due to the hydrophobic nature of ATTO647N as noted by the manufacturer. The solution was covered in foil and left for 2 hours at RT, followed by cleanup using Bio-Spin P-6 desalting columns (Bio-Rad). UV-Vis was used to determine a degree-of-labelling of 0.8. ATTO647N is slightly hydrophobic and we would like to note that higher degree-of-labelling could be achieved with the newer hydrophilic ATTO643 (>2-3), with similar spectral properties.

### Immunolabelling of fixed cells with ATTO647N

1 mL aliquots of Jurkat T cells were transferred from culture flasks into 1.5 mL microcentrifuge tubes and concentrated to 100 µl (*∼* 1-5 million cells/ml) at 300 g for 2 minutes. Cells were incubated in 1% bovine serum albumin (BSA) solution at 4°C for 30 minutes to block non-specific binding. Primary antibody was then added at a given concentration and allowed to bind at 4°C for 50 minutes. For controls (0 pM) no primary antibody was added, but otherwise the same procedure was carried out as for the primary-labelled samples. This was followed by 3x washing in PBS. Cells were then fixed in 2% paraformaldehyde (28906, ThermoFisher) for 20 min at RT, followed by 3x washing in PBS. Cells were kept on ice until flow cytometry or imaging.

### Immunolabelling of live cells with SBB700

1 mL aliquots of Jurkat T cells were transferred from culture flasks into 1.5 mL microcentrifuge tubes and concentrated to 100 *µ*l (*∼* 1-5 million cells/ml) at 300 g for 2 minutes. Cells were incubated in 1% bovine serum albumin (BSA) solution at 4°C for 30 minutes to block non-specific binding. Primary antibody was then added at a given concentration and allowed to bind at 4°C for 50 minutes. For controls (0 pM) no primary antibody was added but otherwise the same procedure was carried out as for the primary-labelled samples. After 3x washing in 1% BSA/PBS, 1 *µ*l of SBB700 secondary antibody was added to the solution and allowed to bind at 4°C for 60 minutes. This empirically determined concentration (unknown proprietary) was higher than manufacturer recommendations. This was followed by 3x washing in PBS. Cells were kept on ice until flow cytometry or imaging.

### Oblique plane microscopy

Oblique Plane Microscopy (OPM) was implemented based on prior work, ^27^ on a custom-built system using a 1.35 NA 100× silicone immersion primary objective lens (Nikon) and a 0.95 NA 40× air immersion secondary objective lens (Nikon), with a custom 358 mm tube lens (ASI), providing a remote refocus magnification of 1.34×. A coverslip was fixed to the front of the secondary objective using an SM1A3 adapter (Thorlabs) with Blu Tack under adjustment until level, confirmed by PSF measurements showing minimal coma. The remote space was imaged through a tertiary glass-tipped objective lens (AMS-AGY v1.0, ASI) and a 200 mm tube lens (TTL200-A, Thorlabs) onto two Photometrics Prime BSI Express sCMOS cameras, one for each channel. The effective pixel size at the sample was 0.116 *µ*m (0.232 *µ*m with 2×2 binning). Imaging at a 30-degree tilt yielded an effective NA of 1.1 (shear-direction) and 1.2 (perpendicular axis). To minimise readout time, both cameras were oriented so that the pixel readout direction was perpendicular to the sample flow.

The laser engine comprised three continuous-wave diode lasers, 405 nm (200 mW, OFL273, Odicforce), 488 nm (55 mW, OFL61, Odicforce), and 638 nm (185 mW, OFL26-1, Odicforce); and a diode-pulsed solid-state laser (561 nm, 500 mW; CNI Lasers), which were combined and coupled into a single-mode fibre (P1-S405-FC-1, Thorlabs) and filtered using a quad-band clean-up filter (ZET405/488/561/640x, Chroma). The output was collimated and shaped into a light sheet using a cylindrical lens (f = 300 mm, LJ1558RM-A, Thorlabs) and an adjustable iris diaphragm was used to adjust the light sheet NA to 0.12. The light sheet was relayed to the sample via an achromatic doublet lens (f = 200 mm, AC254-200-A, Thorlabs) and the primary objective lens. The thickness of the light sheet was determined by imaging the back-reflection from the glass coverslip ^59^ with the beam aligned on-axis (Figure S1). A z-scan was performed to capture the back-reflection intensity across depth, and a Gaussian function was fitted to the line profile to extract the beam waist. The variation of beam width with z-position was then plotted, and the Rayleigh length was calculated from a fit to the data.

Excitation and emission paths were separated by a 3 mm thick quadband dichroic mirror (Di03-R405/488/561/635-t3-25×36, Semrock). After passing the dichroic, the emission was filtered using a quad-band emission filter (ZT405/488/561/640rpcv2, Chroma). After the tertiary objective lens, a longpass dichroic mirror (87-064, Edmund Optics) was used to separate imaging into two channels, one for brightfield imaging and the other one for fluorescence. The fluorescence channel was further filtered using a long-pass emission filter (FELH0650, Thorlabs), while the bright-field used a bandpass filter (FF01-458/64-25, Semrock). An LED (Cree XP-E2 Blue) was used to illuminate the sample with minimal overlap into the fluorescence channel. Oblique illumination was tuned by translating the laser beam toward the periphery of the back focal plane of the primary objective lens. The excitation power density at the sample was 1.12 kW/cm^2^. No scan lenses were required, as the setup used no scanning mirrors. This improved photon transmission compared to most OPM implementations, where pairs of scan lenses are typically used. The transmission efficiency of the OPM path was determined using an alignment laser that was inserted to the primary objective lens from above. The laser power was measured after each optical element resulting in a *∼*58% transmission efficiency for OPM compared to the epifluorescence path (Figure S2).

### Microfluidics

The microfluidic device was fabricated in polydimethylsiloxane (PDMS) with standard rapid prototyping and soft lithography techniques using a Silicon master as a mould as described previously. ^60^ Piranha wet etch (H_2_SO_4_ & H_2_O_2_) was used to clean a 3-inch Silicon wafer, which was then rinsed with deionised water. A 25 *µ*m layer of photoresist SU-8 2025 (Microchem, Warickshire, UK) was coated onto the wafer. Channels were etched into the SU-8 using direct-write laser lithography (MicroWriter ML™, Durham Magneto Optics), using a 375 nm wavelength laser.

PDMS base and cross-linking agent (SYLGARD 184) were mixed at a 1:10 ratio and poured onto the Silicon master, resulting in a negative replica of the SU-8 structures. This was cured for 1 hr at 75°C, forming a hydrophobic elastomer which was peeled away from the master. Device inlet and outlet holes were punched using a Biopsy Puncher, then the PDMS was sealed to a glass slide by treatment with Oxygen plasma. The dimensions of the microfluidic chips were 25 *µ*m height, 400 *µ*m width and 3 cm length.

For smFC experiments, a pipette tip containing 10 *µ*l of labelled cells was attached to the inlet of the microfluidic device. A larger tip containing 100 *µ*l of PBS was placed on top to create a head for gravity flow. An empty pipette tip was attached to the outlet and the flow was initiated using suction created with a pipette. This enabled consistent flow of 5-10 cells/s for OPM imaging at 200 Hz.

### Flow cytometry

Flow cytometry was performed on a CytoFLEX S (Beckman Coulter) controlled by CytExpert software, which was also used for analysis. 20-50 *µ*l of immunolabelled cells were loaded into the instrument’s tube holder. Lasers at 405, 488, 561, and 638 nm were used at their maximum power. Fluorescence was collected through the following channels: 640-660-20 (ATTO647N), 488-690-50 (SBB700) and 561-585-42 (PE). For each acquisition, 10,000-20,000 events were collected. Gating was performed on side (SSC) and forward (FSC) scatter so that only live cells were analysed (and debris were excluded), while doublets were excluded using SSC-A vs SSC-H. Detector gains for each fluorescence channel were set to the maximum. The flow rate was 10-30 *µ*L/min. Quantibrite PE beads (340495, BD Bioscience) were used to quantify the abundance of PE-labelled CD45 on Jurkat T cells.

### Analysis of OPM data

Using the known OPM angle and step size or velocity in the x-direction (typically 400-800 nm/frame) z-stacks were deskewed ^61^ using the deskew function in pycleseperanto. ^62^ Cells were identified by brightfield imaging and the corresponding fluorescence data were manually cropped into individual z-stacks. The deskewed z-stack was then analysed using the single-molecule detection GDSC ^63^ Peak Fit plugin for Fiji. ^64^ Peak Fit parameters for identifying true single-molecule events were determined empirically and remained constant for all analysis. Because the PSF extended over multiple frames in z, resulting in repeat localisations, these were grouped into individual localisations using point cloud cluster segmentation (Matlab’s pcsegdist function). Grouping was done within a distance of 0.6/3 pixels in XY/Z. The threshold for a true single-molecule event was set to a minimum cluster size of 2 points to discard events due to camera pixel noise and cosmic rays.

### Analysis of smFC data

In addition to the processing for OPM data, the smFC pipeline had further steps. Cells were automatically identified by taking a standard deviation z-projection of the deskewed brightfield stacks, followed by Gaussian blurring (20 pixels) and finally the Fiji Find Maxima function. For each identified cell, the intensity was taken as the sum of the background-subtracted (background being a region without a cell) frames in the z-stack. The intensity/molecule calibration was done based on fitting the slope of intensity vs molecule counts on cells within the single-molecule regime (1-35 mol/cell). GDSC Peak Fit was used to find single-molecule events, which were grouped as described above. A manual filter (PSF width < 2 px and SNR > 3) was applied to remove false events, where these parameters were determined empirically. To bridge the single-molecule and high-density regions, we invoked the following conditions:

#### Algorithm 1 Select quantification method

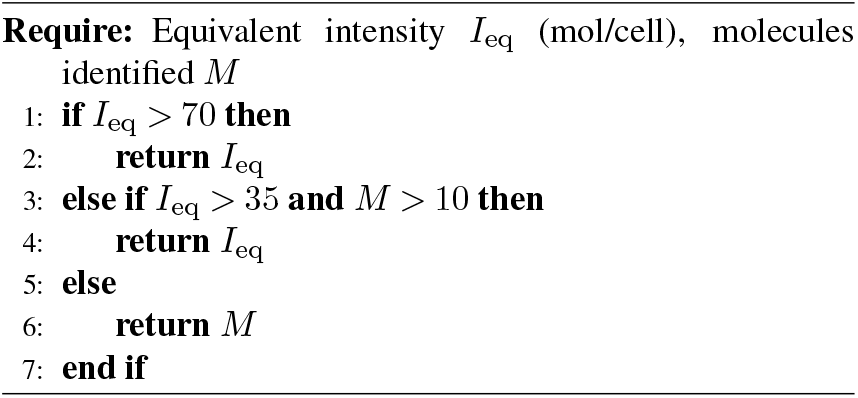

## Notes

### Competing Interest Statement

The authors have declared no competing interest.

