## Supplementary Information for "Single-molecule flow cytometry"

3: Current address: Department of Chemical Engineering and Biotechnology, University of Cambridge, Cambridge, CB3 0AS, UK

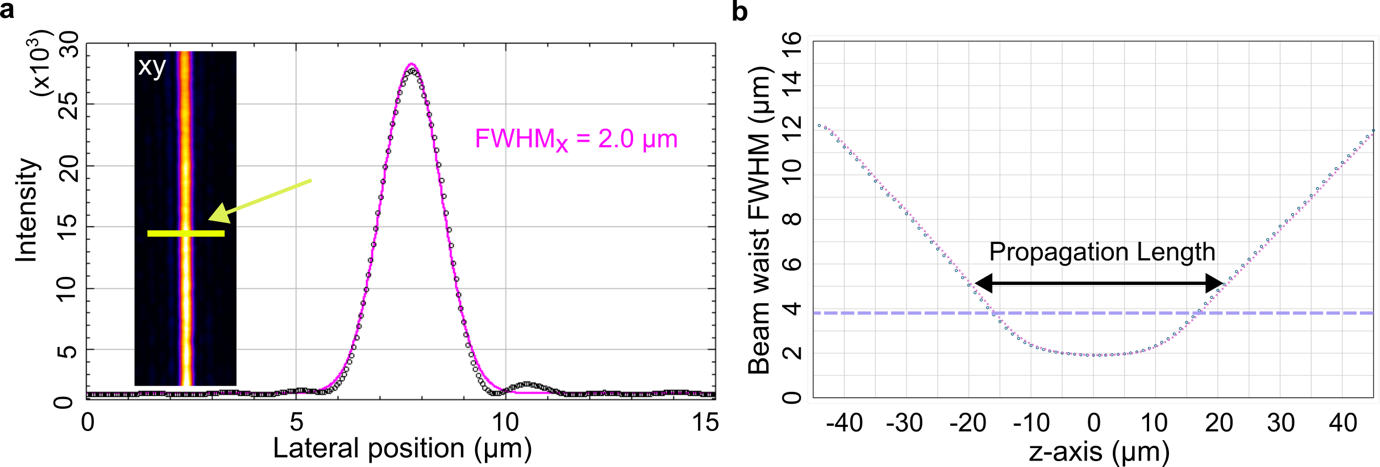

**Figure S1:** (a) Line profile through a cross section of the light sheet to determine its FWHM thickness with a Gaussian fit. Insert is the light-sheet back-reflection from a coverslip. (b) After a z-stack acquisition of the back-reflected light sheet was captured, its thickness at each z-position was determined and was fitted as in (a) to determine its propagation (2×Rayleigh) length.

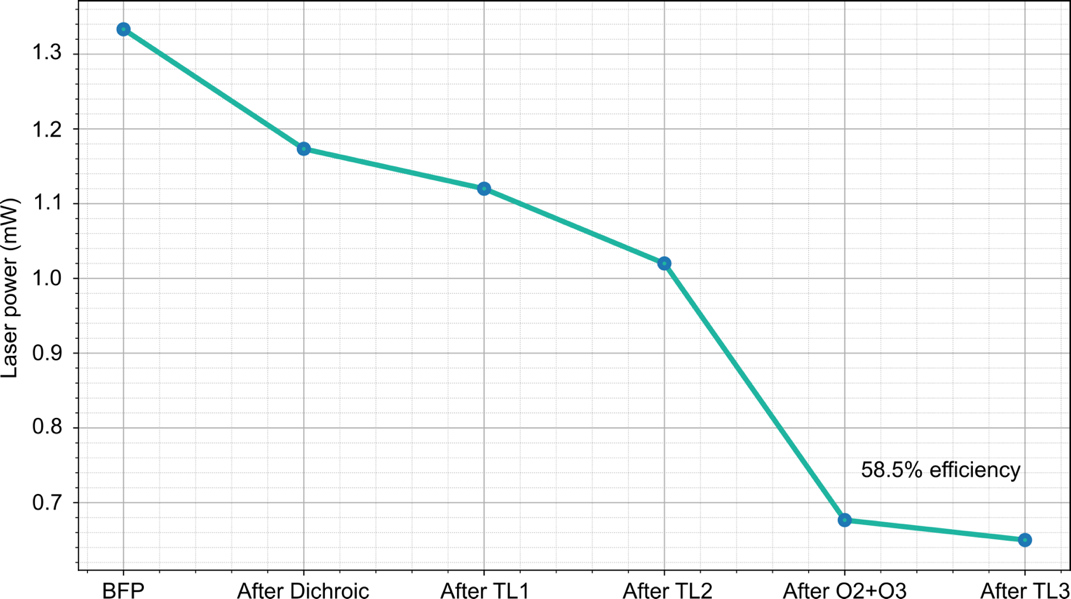

**Figure S2:** The transmission of a laser was evaluated at various positions through the optical train to determine the efficiency of OPM. The efficiency was calculated using “After TL3”/”After TL1”.

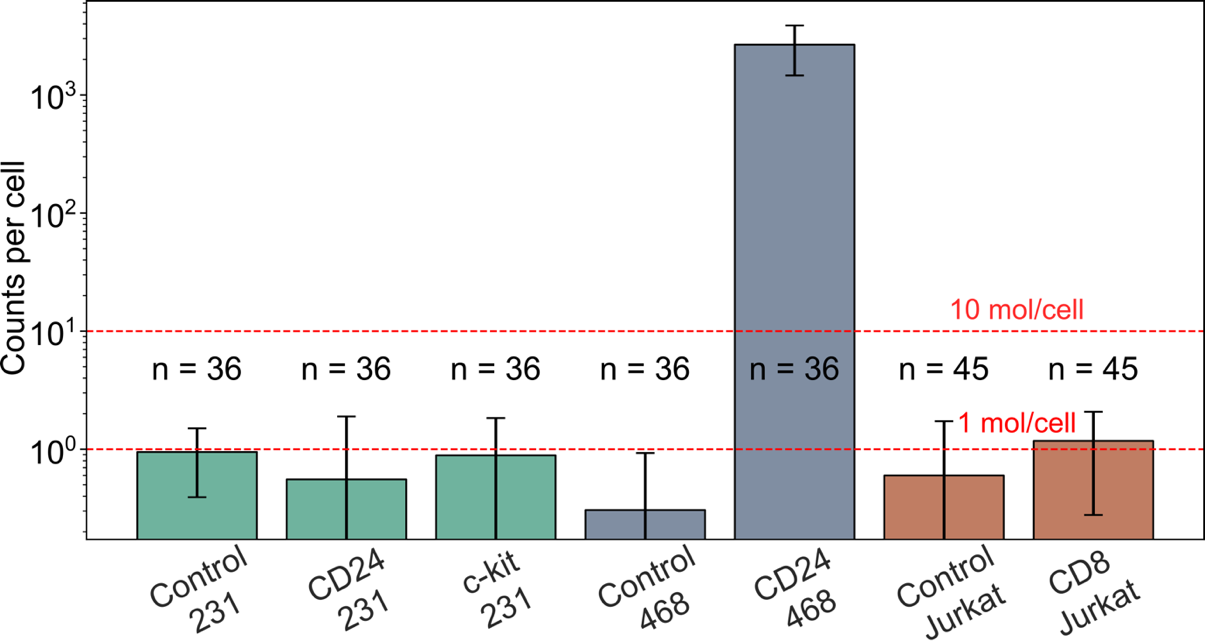

**Figure S3:** Negative (CD24/231, c-kit/231, CD8/Jurkat) and positive (CD24/468) primary antibodies were evaluated using OPM in terms of specificity, demonstrating the ability of smFC to detect 1-10 mol/cell with high confidence. Control refers to no antibody. Reported values are averages and error bars represent standard deviation.

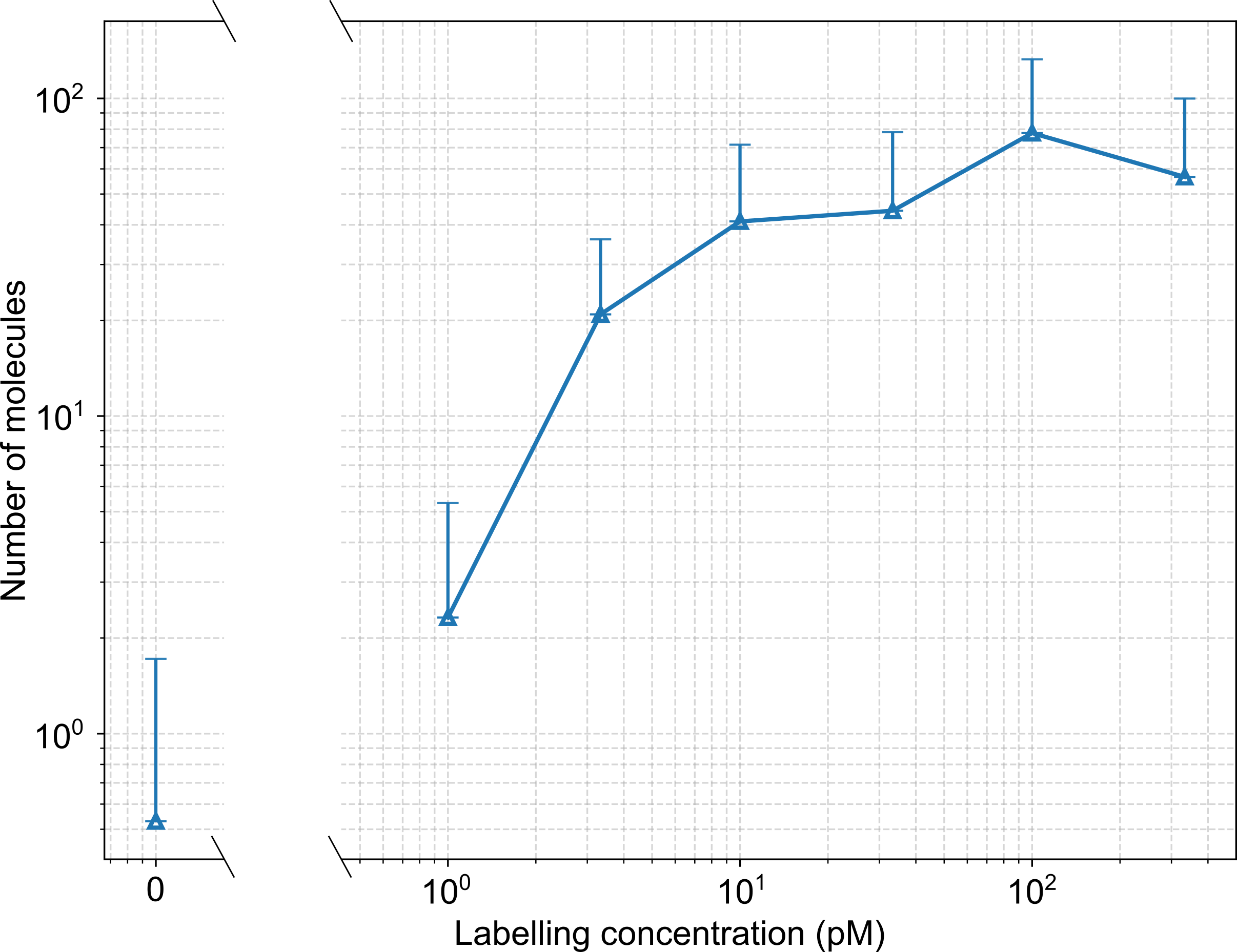

**Figure S4:** The average number of molecules on SBB700/CD45-labelled Jurkat T cells as a function of labelling concentration. Error bars represent standard deviation.
